# Novel non-immunogenic trained immunity inducing small molecule with improved anti-tumor properties

**DOI:** 10.1101/2024.03.22.585780

**Authors:** Jainu Ajit, Hannah Riley Knight, Qing Chen, Ani Solanki, Jingjing Shen, Aaron P. Esser Kahn

## Abstract

Trained immunity refers to the non-specific innate immune memory response triggered by the epigenetic and metabolic rewiring of innate immune cells. A strengthened innate immune system significantly improves disease resistance. However, very few trained immunity-inducing molecules have been identified. Almost all molecules for training are primarily immunogenic and then subsequently induce training. Non-immunogenic molecules that induce training could be employed in therapies without the concern of adverse inflammatory reactions. We identified a small molecule, A1155463, that modulates cellular metabolism to induce trained immunity in macrophages *in-vitro*. We show that nanomolar concentrations of these compounds uniquely alter only cellular metabolism without leading to apoptosis. We further observed that these compounds could induce training in an *in-vivo* model in mice. A1155463 training improved anti-tumor resistance to B16.F10 melanoma cells. The effect was enhanced upon combination with checkpoint therapy. In summary, we report the discovery of a novel trained immunity-inducing small molecule with enhanced anti-tumor properties.

## Introduction

Innate immune cells form the first line of defense and protect against disease.^1^ Innate immune cells initiate their activity by responding to pathogen and damage-associated molecular patterns (PAMPs, DAMPs).^2,3^ In addition to inflammatory immune activation, certain stimuli induce a so-called non-specific memory response called trained immunity.^4^ These stimuli induce epigenetic and metabolic changes in the cell, which facilitate increased transcription and translation of pro-inflammatory cytokines following subsequent activation of the cells.^5^

To date, inducers of trained immunity have predominantly been activators of innate immunity including, vaccines such as BCG; infections: candidiasis, malaria;^6–9^ cell wall component, β-glucan, muramyl dipeptide (MDP);^10,11^ TLR agonists, including CpG (TLR 9) and flagellin (TLR 5);^12,13^ danger signals, including heme and oxidized LDL;^14–16^ and hormones involved in immunity, including aldosterone and catecholamines.^17,18^ Most trained immunity inducers alter glycolysis or oxidative phosphorylation.– promoting the faster generation of ATP for activation when the cells are challenged later.^19-21^ Indeed, both exogenous fumarate and mevalonate also induce training.^19,20^ Increased production of metabolites like acetyl-CoA modulate the trained cell’s epigenetics promoting faster transcription and translation of pro-inflammatory genes following disease challenge.

As a result of the metabolic and epigenetic changes, a trained cell can better react to a secondary trigger. Although trained immunity has beneficial effects, such as better anti-tumor responses in mice, uncontrolled activation and training by many of the listed molecules can be detrimental.^21^ Extensive studies have reported the role of trained immunity in allograft rejection, allergy, neurodegenerative disorders, and atherosclerosis.^22–27^ To harness the beneficial effects of training, there is a need to identify novel molecules that are both non-immunogenic and allow yet induce responses analogous to current training compounds.

We report a novel non-immunogenic trained immunity inducing molecule-A1155463-an inhibitor of the anti-apoptotic Bcl-xL protein.^28^ Bone-marrow derived macrophages (BMDMs) trained with A1155463 at nanomolar concentrations secreted higher pro-inflammatory cytokines following an LPS challenge compared to untrained cells. Furthermore, we observed that the training mechanism depended on glycolysis as inhibition with 2-deoxy glucose removed training effects observed *in-vitro*. A1155463 also improved training effects in an *in-vivo* model in mice and improved tumor resistance. The anti-tumor effects were further enhanced when combined with checkpoint blockade, demonstrating the potential to use trained immunity to enhance current therapeutics.

## Results and discussion

B-cell lymphoma 2 (BCL-2) family proteins regulate programmed cell death or apoptosis.^29^ Small molecule inhibitors targeting single or multiple anti-apoptotic Bcl-2 proteins have been developed to induce apoptosis for anti-cancer therapeutics.^30^ In addition to regulating apoptosis, Bcl-2 family proteins also modulate cellular metabolism.^31–34^ As changes in cellular metabolism can induce trained immunity, we hypothesized that inhibiting anti-apoptotic Bcl-2 proteins can cause training.^35^ Within the Bcl-2 family of proteins, B-cell lymphoma-extra-large (Bcl-xL) is an anti-apoptotic protein in the mitochondria.^36^ Several inhibitors of Bcl-xL protein exist and a few have been reported to exhibit undetermined immunomodulatory effects. Of these, A1155463, inhibits Bcl-xL and alters immune responses though the mechanism has never been documented.^28,37^

To test if A115463 induced training, we first performed an *in-vitro* training assay in BMDMs (**Fig. 1a**).In this assay, BMDMs were trained with either PBS (untrained or control group) or A1155463 at various concentrations for 24 h. After 24 h, the cells were washed, and the media was replaced. The cells were allowed to rest for four days. On day 6, cells were washed and challenged with Pam3 (TLR 1/2 agonist). The supernatant was tested 24 h following the challenge. Trained cells produce a higher concentration of pro-inflammatory cytokines, namely. IL-6 and TNF-α, than untrained cells following challenge with a TLR agonist. In line with our hypothesis, we observed an approximately 1.2-fold increase in both IL-6 and TNF-α following the challenge with Pam3 (**Fig. 1b**).

**Fig. 1:**
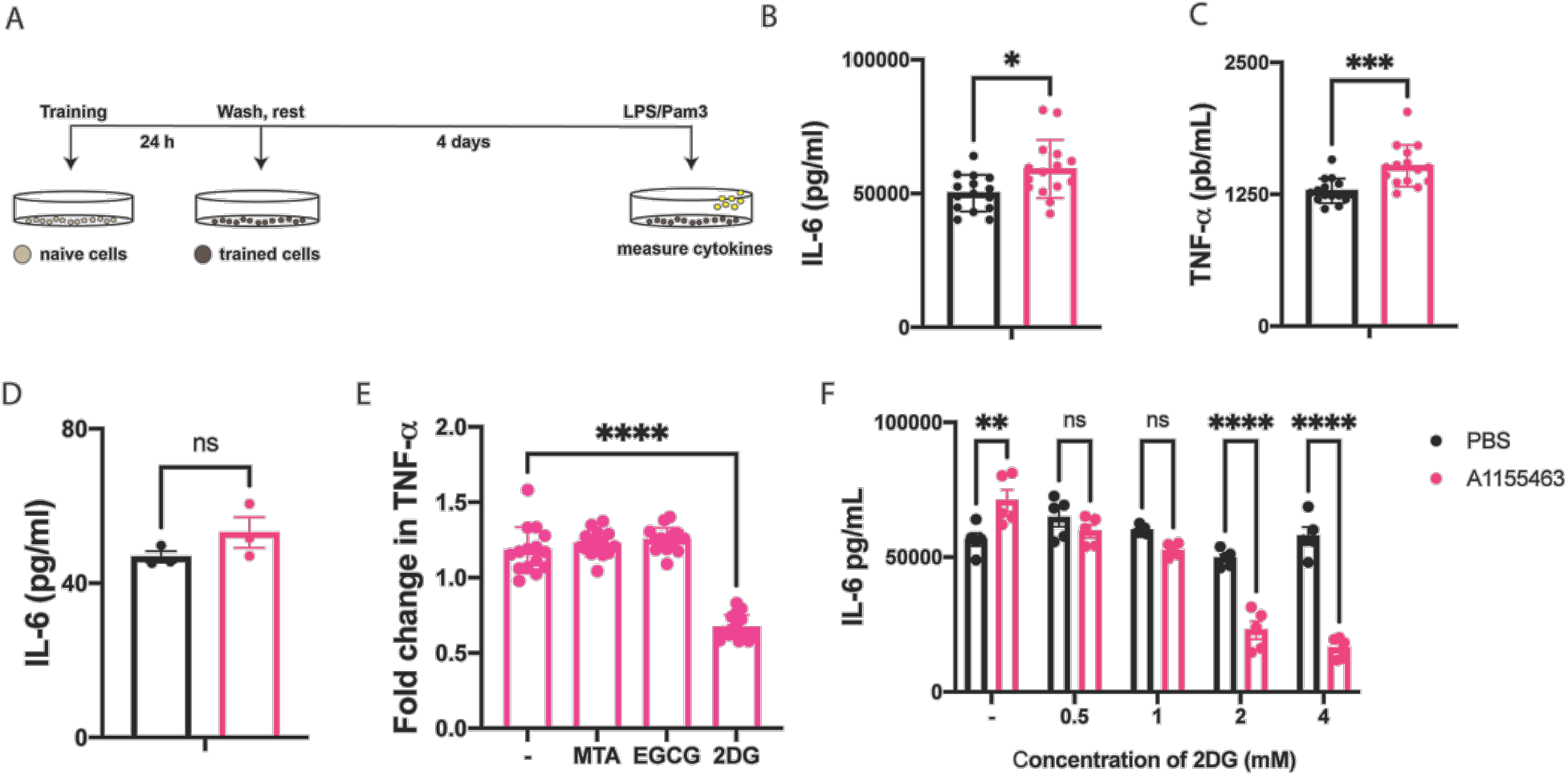
Testing A1155463 as an inducer of TI in vitro. a) Schematic for in-vitro TI assay in BMDMs. b) IL-6 levels 24 h post Pam3 challenge in an in-vitro TI assay in BMDMs trained with either PBS (black) or A1155463 (pink); n = 15 c) TNF-α levels 24 h post Pam3 challenge in an in-vitro TI assay in BMDMs trained with either PBS (black) or A1155463 (pink); n = 10 d) IL-6 levels 24 h post-stimulation with PBS (black) or A1155463 (pink); n = 3 e) Fold change in TNF-α following Pam3 challenge in BMDMs trained with A1155463 pre-treated with PBS, MTA (histone methyl transferase inhibitor), EGCG (histone acetyltransferase inhibitor) or 2-DG (glycolysis inhibitor); n = 15, significance compared with PBS control of each inhibitor. f) IL-6 levels 24 h post Pam3 challenge in an in-vitro TI assay in BMDMs trained with either PBS (black) or A1155463 (pink) pretreated with varying concentrations of 2-DG; n= 5. All values are expressed as mean ± SEM, and statistics were conducted using unpaired student’s T-test or 2-way ANOVA with Sidak’s multiple comparisons test (significance compared with PBS group). \**P* < 0.05, \*\**P* < 0.01, \*\*\**P* < 0.001, \*\*\*\**P* < 0.0001, n.s., not significant.

Bcl-xL is an anti-apoptotic protein with some role in mediating inflammatory reactions.^38,39^ Most documented inducers of trained immunity elicit an initial immune response upon administration. However, we conjectured that A1155463 might be different as it had never been reported elicit inflammatory responses in vitro. To test if A1155463 elicited an enhanced innate response on its own in vitro, supernatant from BMDMs cultured with A1155463 for 24 h was analyzed. We observed no significant difference in IL-6 levels between PBS-treated and A1155463-treated BMDMs after 24 h (**Fig. 1c**). We also screened for various other cytokines 24 h post treatment and observed significantly low levels (below 100 pg/mL) in the cell supernatant. A1155463-treated cells consistently produced low levels of MCP-1(around 70 pg/mL) irrespective of concentration which has been previously described in macrophages.^40,41^ This is noteworthy as a typical inflammation, induced by a TLR 1/2 agonist like Pam3, induces 20 times higher MCP-1 levels (around 1500 pg/mL) in BMDMs. Furthermore, we validated the non-immunogenicity of A1155463 using an NF-κb reporter cell line. We observed no significant difference between PBS-treated and A1155463-treated RAW-Blue™ after 18 h (**Fig. SI 2**). Taken together, this evidence supports that A1155463 is a non-immunogenic trained immunity inducing molecule.

Since Bcl-xL is a known anti-apoptotic protein, we next tested to see if inhibition with 100 nM of A1155463 induced apoptosis in BMDMs. To test this, we used Annexin-V staining to gate for apoptotic cells 24 h following stimulation with either PBS or A1155463. We observed no significant difference in the percentage of apoptotic cells in A1155463 treated cells (**Fig. SI 3**). From this, we conclude that A1155463-induced training does not operate through an apoptotic mechanism.

After ruling out apoptosis and innate inflammation, we were interested in determining the mechanism of action of A1155463-induced training in BMDMs. Training induces epigenetic and metabolic rewiring of macrophages which can be probed using small molecule inhibitors of various pathways.^42^ Epigenetic changes are associated with increased histone methylation and acetylation, promoting transcription and translation of pro-inflammatory cytokines in response to pathogenic stimuli. Metabolic changes alter cellular metabolism, mostly favoring glycolysis, to meet the higher energy requirements of a trained cell. In this assay, BMDMs were pre-treated with small molecule inhibitors, namely, MTA (histone methyl transferase inhibitor), EGCG (histone acetyltransferase inhibitor), or 2-deoxy glucose or 2-DG (glycolysis inhibitor) for 30 min. This was followed by the standard training assay as described before (**Fig. 1a**). For epigenetic small molecule inhibitors we observed no significant difference. However with the glycolysis inhibitor, 2DG pre-treatment caused a 50% reduction in the TNF-a level following Pam3 challenge (**Fig. 1d**). We also noted a significant decrease in Pam3 induced IL-6 levels for different concentrations of 2-DG. (**Fig. 1e**) This result indicates that A1155463 induced training likely proceeded through metabolic changes in BMDMs. This matches the previous literature showing Bcl family proteins modulate cellular metabolism.^31–34^ Of note, to observe this inhibition of glycolysis, it was necessary to increase the concentration of 2DG from 1 mM to between 2 and 4 mM. This result suggests that glycolytic inhibition and therefore glycolysis may vary between inducers of trained immunity. (**Fig. 1f**) Trained immunity is typically associated with either epigenetic or metabolic alterations, and these mechanisms often influence each other. While many recognized training molecules primarily regulate epigenetic processes, our molecule is among the few that primarily induces metabolic changes. Notably, these metabolic alterations don’t result in immediate but rather gradual epigenetic changes, a phenomenon not measured in our experimental protocol. The concept of metabolism governing downstream epigenetic modifications has been documented for molecules such as BCG.^43^ In essence, we aim to emphasize that our molecule’s primary mode of action is through inducing metabolic changes. The subsequent training effects may involve both epigenetic and metabolic alterations, but the intricacies of this interplay cannot be fully elucidated within the confines of our current experimental design, especially considering the limitations imposed by the use of small molecule inhibitors.

Since A1155463 induced training effects in-vitro, we next tested its ability to induce training in mice. The half-life of Bcl-xL inhibitors in mice was previously determined to be around 9 h.^28^ To ensure sufficient material for training in-vivo, we used a dosage of 15 mg/kg administered intra-peritoneally daily for 6 days (**Fig. 2a**). Vehicle control group was also dosed similarly to account for solvent effects. To rule out the impact of innate immune priming due to the high dosage, we first checked for systemic cytokines 24 h after the last dosing on day 6. We observed no systemic IL-6 or TNF-a, indicating no priming or adverse inflammatory reaction to the compound (**Fig. SI 4**). A separate set of trained mice were challenged with LPS (5 μg per mouse), and systemic cytokines were analyzed after 1 h. We observed 2.3-fold and 1.97-fold increase in IL-6 and TNF-a levels respectively following LPS challenge suggesting enhanced in-vivo training effects. (**Fig. 2b**).

**Fig. 2:**
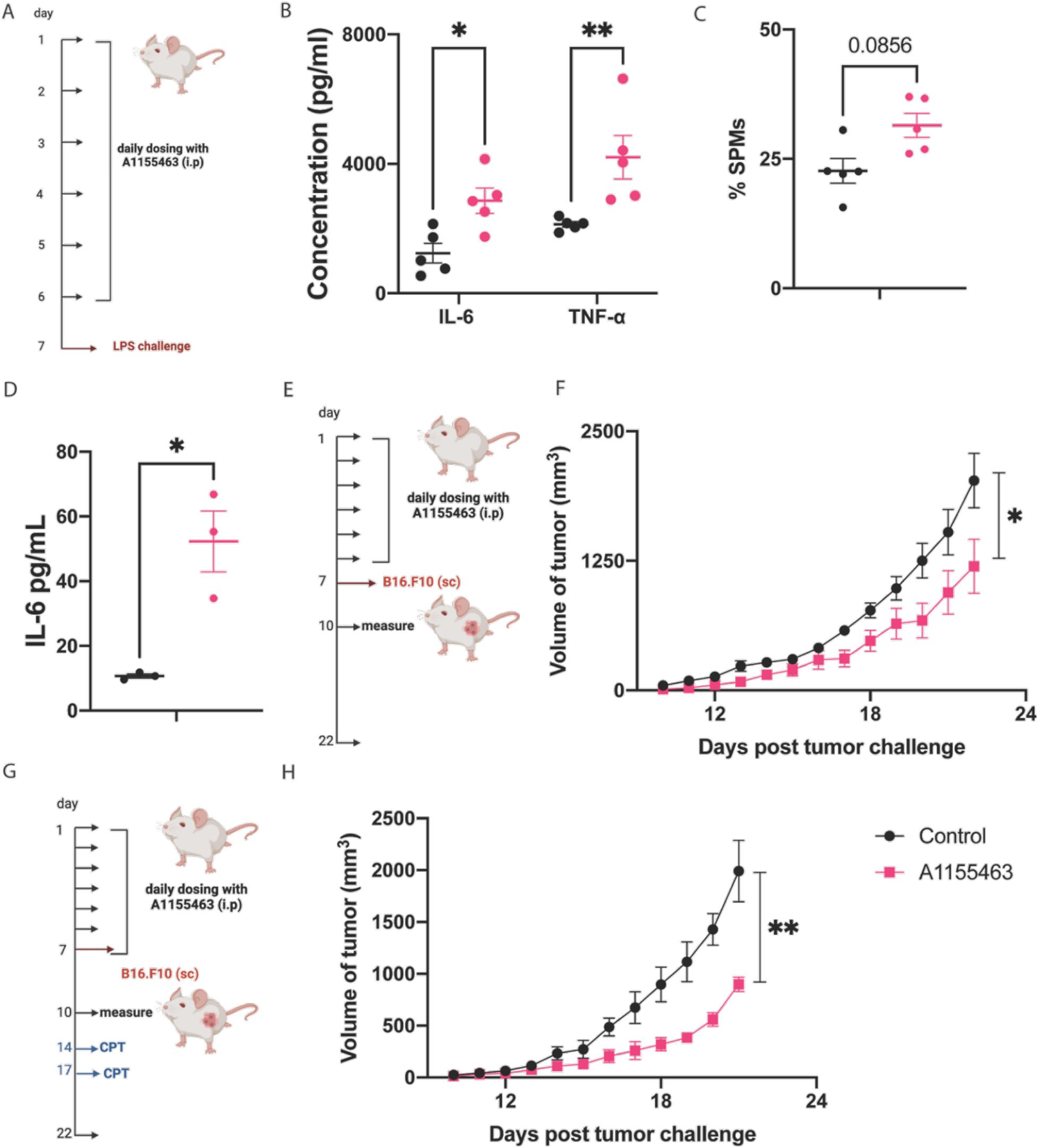
in-vivo TI assays. a) in-vivo training schedule-mice were dosed i.p. with training material (PBS or A1155463 at 15 mg/kg) daily for six days. Mice were challenged with 5 μg LPS on day 7. b) IL-6 and TNF-α levels 1 h following LPS challenge with either PBS (black) or A1155463 (pink); n= 5. c) % of SPMs from the peritoneal cavity of PBS (black) or A1155463 trained (pink) mice 2 h post LPS challenge on day 7. d) IL-6 levels from bone-marrow cells isolated on day 7 challenged ex-vivo with LPS from either PBS (black) or A1155463 (pink) trained mice; n= 3 e) in-vivo training schedule for B16.F10 tumor challenge f) Tumor volumes of mice trained with PBS (black) or A115463 (pink) mice; n= 8. g) in-vivo training schedule for B16.F10 tumor challenge with checkpoint therapy (CPT) administered on day 14 and 17. h) Tumor volumes of mice trained with PBS (black; n=4) or A115463 (pink; n=5).

Apart from increased systemic cytokines, recruitment of monocyte-derived small peritoneal macrophages (SPMs) is another hallmark of enhanced training. We observed a non-significant increase in the percentage of SPMs two hours post LPS challenge (**Fig. 2c**). Central effects of trained immunity occur at the bone-marrow progenitor favoring myelopoiesis.^44^ To test if A1155463 induced training effects in bone-marrow cells, we isolated cells from the bone-marrow seven days following the training schedule described before (**Fig. 2a**). We challenged bone-marrow cells from PBS or A1155463 trained cells with 5 ng/mL of LPS and analyzed the supernatant 24 h later. We observed an approximately 5-fold increase in bone-marrow derived IL-6 levels from A115463 trained mice than untrained control (**Fig. 2d**). Taken together, these results indicate enhanced in-vivo training effects of A115463.

To demonstrate the clinical translatability of A115463-derived training effects, we next turned to a B16.F10 tumor model. In this model, mice were trained with either vehicle control or A1155463 (15 mg/kg) daily for 6 days. Then, 24 h post the last dosing, mice were challenged subcutaneously with B16.F10 tumors (**Fig. 2e**). We observed a significant decrease in tumor growth in A1155463 trained mice compared to the untrained (vehicle) control group (**Fig. 2f**). Encouraged by these results, we next tested to see if A1155463 training can be combined with conventional checkpoint therapy to improve anti-tumor effects further.^44^ We followed the same schedule as before and added two doses of checkpoint inhibitors - anti-PDL1 and anti-CTLA-4 combination on days 14 and 17 after the first training regimen (**Fig. 2g**). We observed significantly reduced tumor growth in A1155463 trained mice that were supplemented with checkpoint therapy (**Fig. 2h**). Taken together, these results demonstrate that A1155463 mediated training leads to enhanced anti-tumor effects. Previous studies have identified macrophage polarization into pro-inflammatory M1 phenotype following inhibition of Bcl2 and Bcl-xL inhibition in tumor models.^45–47^ We hypothesize that A115463-induced training similarly results in the reprogramming of tumor associated macrophages into M1 phenotype contributing to the observed anti-tumor effects. However, further studies need to be performed to dissect these cellular players and reveal novel drug delivery approaches in developing anti-cancer therapeutics.

## Conclusions

Trained immunity improves our first line of defense – innate immune cells.^4^ Very few, small molecule inducers of trained immunity have been identified thus far. The most well studied training molecules include BCG vaccine and β-glucan, both of which are immunogenic, activating NOD-2 and Dectin-1 receptors, respectively.^6-10^ Other identified molecules are primarily based on the metabolic and epigenetic pathways that mediate training. To date, A1155463 is distinctive from these other molecules in that (a) it is a drug-like molecule and (b) it does not elicit inflammation upon initial administration. The second point in particular may be quite valuable for therapeutic applications.

Identifying new classes of novel non-immunogenic molecules that elicit training can help to harness the beneficial effects of training by increasing available dosing regimens and routes of administration. We hypothesized that targeting other proteins involved in cellular metabolism would present a fruitful approach to identifying novel training compounds. The Bcl-2 family of proteins, due to its location on the mitochondrial membrane, are known regulators of cellular metabolism. Our hypothesis was validated in that molecule inhibitor, A1155463 targeting Bcl-xL, induced trained immunity both in vitro and in vivo.

We observed a significant increase in training effects in A1155463-trained BMDMs compared to untrained controls. We noted that, indeed, this effect was mediated by an increase in glycolysis by inhibitor assays. We then tested for A1155463-mediated training effects in-vivo and noticed enhanced pro-inflammatory cytokines following LPS challenge compared to untrained mice. This result encouraged us to test for training-induced anti-tumor capabilities using a B16.F10 model. We observed that A1155463-mediated training improved anti-tumor effects by itself and in combination with checkpoint blockade. In summary, we report a novel non-immunogenic training immunity inducing small molecule with immuno-modulatory potential. Future studies must decipher the cellular players involved in this training pathway to achieve controlled and safe training effects.

## Methods

All chemicals and reagents unless noted otherwise were purchased from Sigma. ELISA kits and all fluorescently tagged antibodies were purchased from BioLegend. Pam3, LPS and OVA were purchased from Invivogen. All cell culture reagents were obtained from Thermo Fisher Scientific. Cells were maintained at 37 °C and 5% CO2. C57Bl/6J mice were obtained from Jackson Laboratories and acclimatized for 1 week prior to experimentation. All animal experiments were conducted with approval from the University of Chicago Institutional Animal Care and Use Committee (approval number 72517). All statistical analyses were performed using GraphPad Prism.

### *In-vitro* training assay

Monocytes were harvested from the femurs of 6 week old C57BL/6 mice (Jackson Laboratory). Monocytes were differentiated into macrophages using supplemented culture medium: RPMI 1640 (Life Technologies), 10% heat inactivated fetal bovine serum (HIFBS) 2 × 10^-3^ m L-glutamine (Life Technologies), antibiotic antimycotic (1×) (Life Technologies), and 10% MCSF (mycoplasma free L929 supernatant) for 5 d at 37 °C and 5% CO_2_. The cells were then released with 5 × 10^-3^ m EDTA in PBS, counted and plated at desired densities. BMDMs were plated at a density of 100,000 cells per well in flat bottom 96-well plates (Corning) at a final volume of 200 µL) and rested for a few hours to adhere at 37 °C and 5% CO_2_. After the cells were adherent, training material was added at the desired concentration and incubated for 24 h. Cell supernatant was collected after 24 h and stored at -20 degrees for cytokine analysis. Then cells were washed and rested for 3 d. On day 4, BMDMs were washed again and primed with 25 ng mL^-1^ IFN-*γ* (BD Biosciences) for 24 h. On day 5, a final wash was performed, and cells were stimulated with 10 ng mL^-1^ Pam3CSK4 (Invivogen). Cell supernatant was collected after 24 h to measure IL-6 and TNF-*α* levels using ELISA (BioLegend) according to manufacturer’s instructions.

#### Epigenetic and metabolic inhibition assays

For inhibition studies, BMDMs were preincubated for 30 min with inhibitors before stimulating with training materials and were left in the media during the 24 h training period.^[23]^ For epigenetic pathway analysis, cells were pretreated with 500 uM 5′-deoxy-5′-(methylthio)adenosine (MTA), 6 uM pargyline and 50 uM (-)-epigallocatechin-3-gallate (EGCG) (all from Sigma Aldrich). For metabolic pathway analysis, cells were pretreated with indicated concentrations of deoxy glucose (Sigma). This was followed by the regular training regimen as described earlier.

#### RAW-Blue NF-κB assay

RAW-Blue NF-κB cells (InvivoGen) were passaged and plated in a 96-well plate at 100,000 cells per well in 180 μl of Dulbecco’s modified Eagle’s medium (DMEM) containing 10% heat-inactivated fetal bovine serum (HIFBS). Cells were incubated at 37°C and 5% CO_2_ for 1 hour. PBS or A1155463 was added at indicated concentrations; LPS was used at a concentration of 100 ng/mL as a positive control, the volume of each well was brought to 200 μl and incubated at 37°C and 5% CO_2_ for 18 hours. After 18 hours, 20 μl of the cell supernatant was placed in 180 μl of freshly prepared QuantiBlue (InvivoGen) solution and incubated at 37°C and 5% CO_2_ for up to 2 hours. The plate was analyzed every hour using a Multiskan FC plate reader (Thermo Fisher Scientific), and absorbance was measured at 620 nm.

#### Apoptosis assay

10^6 BMDMs were plated in 12 well plates and stimulated with PBS, A1155463 (100 nM) or camptothecin (2 mM) in a total volume of 1 mL for 24 h at 37°C and 5% CO_2_. After 24 h, cells were washed with cold PBS, resuspended in binding buffer and stained with 5 uL of FITC-AnnexinVfor 15 min in the dark according to manufacturer’s protocol (BD Biosciences-FITC annexin V apoptosis detection kit). Cells were analyzed for flow cytometry to detect apoptotic cells.

#### in-vivo training assay

Mice were trained intraperitoneally daily for 6 days (ip) with vehicle control or 15 mg/kg A1155463. One set of mice were subjected to retro-orbital bleeding 24 h post last training dose (day 7) and cytokines were analyzed for innate immune priming effects. Bone marrow was isolated from these mice and analysed ex-vivo. Briefly, 100,000 bone marrow cells were plated in 96 well plates and challenged with 5 ng/mL LPS. Cell supernatant was analysed using IL-6 ELISA kits to measure central training effects. A separate set of mice were challenged intraperitoneally with 5 µg LPS (serotype O55:B5; Invivogen) serum was collected after 1 h. Serum cytokines were analyzed using Legendplex Mouse Inflammation Panel (BioLegend) according to the manufacturer’s protocol.

#### Small peritoneal macrophage analysis

Mice were subject to the same training schedule as described above. 2 h after LPS challenge, mice were euthanized and peritoneal cells were harvested by lavage. Cells were analysed by flow cytometry to determine the percentage of small peritoneal macrophages (CD11b+ MHC2hi F4/80-). All antibodies were purchased from BioLegend or BD Biosciences (anti-mouse phycoerythrin (PE) CD11b, APC-MHC2, Pe/Cy7-F4/80)

#### Tumor challenge

Mice were anesthetized and shaved of hair from their right-side using clippers. Following grooming, the mice were injected with 200,000 B16F10 melanoma in 50 mL of PBS subcutaneously. Mice were first trained with indicated materials following training regimen described above. 24 h post last training, all the animals were challenged with B16.F10 tumor. Tumor progression was monitored and measured with a caliper, recording the width, length, and height of the tumor every other day throughout the experiment. Mice with tumor sizes exceeding 20 mm along any direction were sacrificed. For checkpoint therapy, mice were injected ip with 100 ug each of anti PDL-1 and anti-CTLA-4 twice on days 14 and 17 post tumor challenge. Tumors were measured until endpoint.

#### Statistical Analysis

All values are expressed as mean ± SEM. The sample size is as indicated in figure captions in all *in-vivo* and *in-vitro* experiments. Student’s T-test was applied for comparing two groups, and one- or two-way analysis of variance (ANOVA) followed by Tukey’s or Dunnett’s multiple comparisons for comparison of multiple groups (indicated in the figure captions) using the GraphPad Prism 9 software. *P* values less than 0.05 were considered statistically significant. Significance \**P* < 0.05, \*\**P* < 0.01, \*\*\**P* < 0.001, \*\*\*\**P* < 0.0001, n.s., not significant.

## Acknowledgement

The authors thank University of Chicago veterinary technicians for exceptional animal care.

