## Supplemental Information for "Novel non-immunogenic trained immunity inducing small molecule with improved anti-tumor properties"

### 1.1 Supplementary Figures

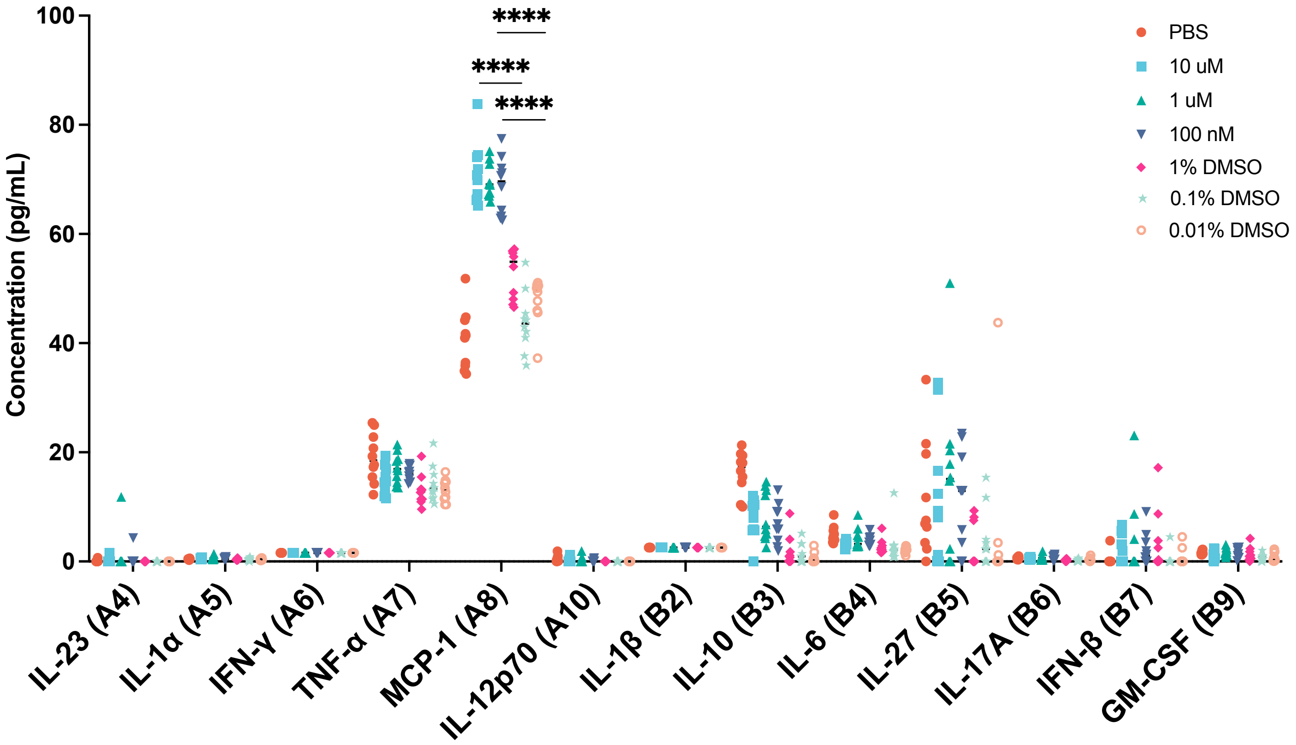


**Supplementary Figure 1.** Testing immunogenicity of A115463 on BMDMs: BMDMs (100,000 cells per well) were incubated with PBS (red), different doses of A1155463 (blue 10 uM, green 1 uM or purple 0.1 uM) or different concentrations of DMSO, reflective of the A1155463 doses for 24 h. Supernatant was tested using a Legend plex mouse inflammation kit; n= 10; All values are expressed as mean ± SEM, and statistics were conducted using two-way ANOVA with Tukey's multiple comparisons test (significance compared as indicated). **P* < 0.05, ***P* < 0.01, ****P* < 0.001, *****P* < 0.0001, n.s., not significant.

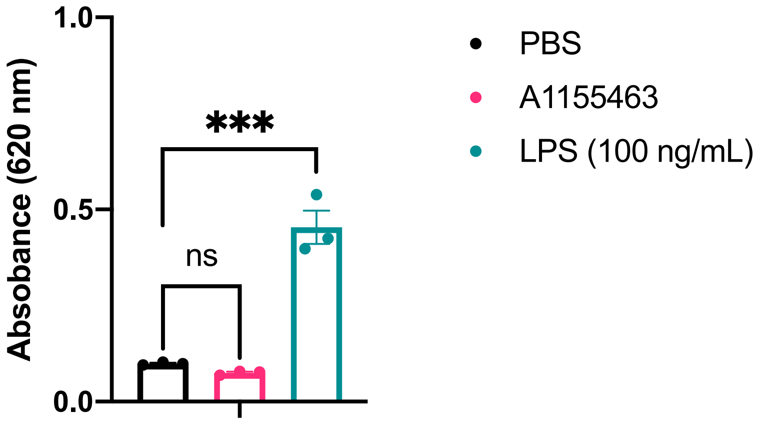


**Supplementary Figure 2.** RAW-Blue assay: RAW-Blue cells (100,000 cells per well) were incubated with PBS (black), A1155463 (pink) or 100 ng/mL LPS (green) for 18 h. Supernatant was tested using Quanti-Blue assay and absorbance measured at 620 nm; n= 3; All values are expressed as mean ± SEM, and statistics were conducted using one-way ANOVA with Dunnett's multiple comparisons test (significance compared with PBS group). **P* < 0.05, ***P* < 0.01, ****P* < 0.001, *****P* < 0.0001, n.s., not significant.

1) unstained control 2) PBS control 3) positive control: camptothecin

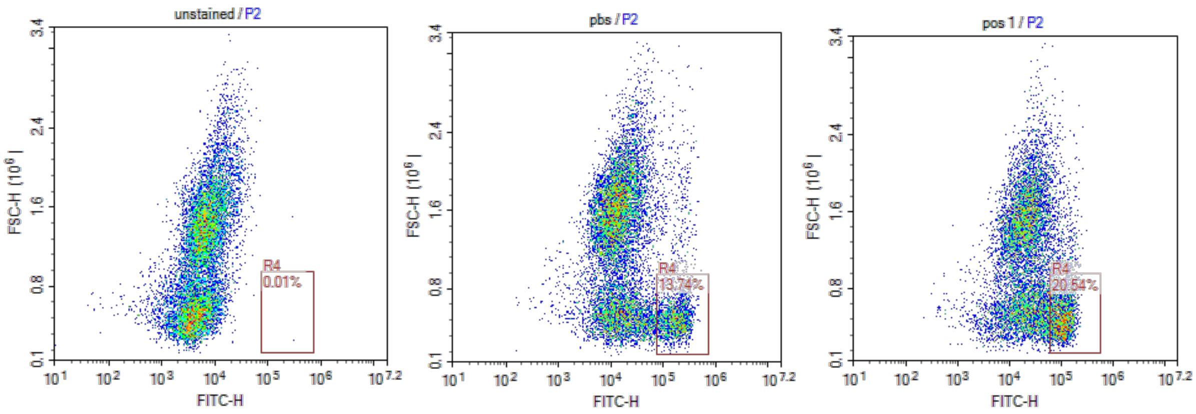


Representative flow plot for A115463 at 100 nM

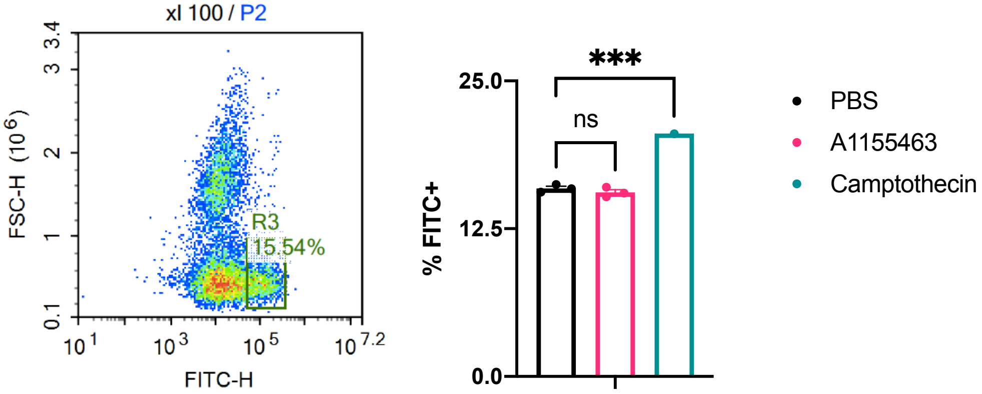


**Supplementary Figure 3.** Representative plots for Annexin V gating strategy for apoptotic cells: BMDMs were stimulated with PBS (negative control), Camptothecin (positive control) or A1155463; n = 3, (except camptothecin n=1), statistics were conducted using one-way ANOVA with Dunnett's multiple comparisons test (significance compared with PBS group). *$P < 0.05$, **$P < 0.01$, ***$P < 0.001$, ****$P < 0.0001$, n.s., not significant

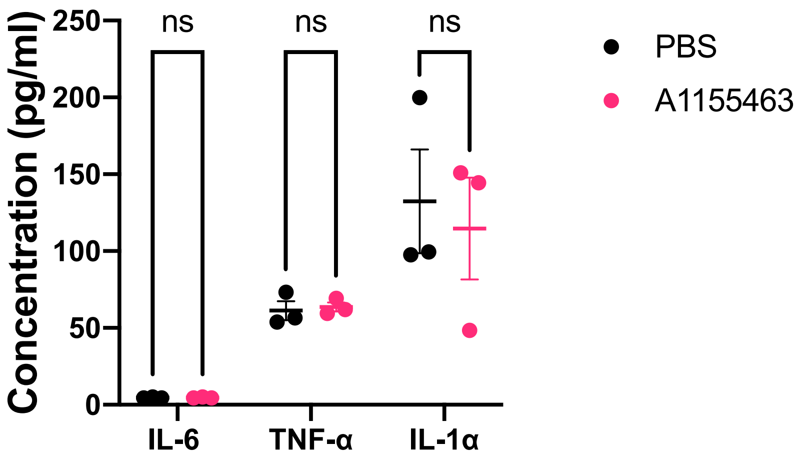


**Supplementary Figure 4.** Ruling out innate immune priming: Mice were trained with PBS or vehicle (black) or A1155463 (pink) according to schedule illustrated in Fig. 2a. 24 h after the last dosing, mice were bled and serum analyzed for pro-inflammatory cytokines using cytokine bead array; n= 3; statistics were conducted using one-way ANOVA with Sidak's multiple comparisons test (significance compared with PBS group). *$P < 0.05$, **$P < 0.01$, ***$P < 0.001$, ****$P < 0.0001$, n.s., not significant
